# Heliconiini butterflies can learn time-dependent reward associations

**DOI:** 10.1101/2020.06.06.135459

**Authors:** M. Wyatt Toure, Fletcher J. Young, W. Owen McMillan, Stephen H. Montgomery

## Abstract

For many pollinators, flowers provide predictable temporal schedules of resource availability, meaning an ability to learn time-dependent information could be widely beneficial. However, this ability has only been demonstrated in a handful of species. Observational studies of *Heliconius* butterflies suggest that they may have an ability to form time-dependent foraging preferences. *Heliconius* are unique among butterflies in actively collecting and digesting pollen, a dietary behaviour linked to spatiotemporally faithful ‘trap-line’ foraging. Time-dependency of foraging preferences is hypothesised to allow *Heliconius* to exploit temporal predictability in alternative pollen resources, as well as contributing to optimal use of learnt foraging routes. Here, we provide the first experimental evidence in support of this hypothesis, demonstrating that *Heliconius hecale* can learn opposing colour preferences in two time periods. This shift in preference is robust to the order of presentation, suggesting that preference is tied to the time of day and not due to ordinal learning. However, we also show that this ability is not limited to *Heliconius*, as previously hypothesised, but is also present in a related genus of non-pollen feeding butterflies. This demonstrates that time learning pre-dates the origin of pollen-feeding and may be prevalent across butterflies with less specialized foraging behaviours.

## 1. Introduction

The specific cues providing animals with reliable information about resource availability vary across environments, and this variation shapes which cues animals learn to use [1,2]. For example, animals inhabiting dark environments may preferentially learn associations with olfactory cues over visual cues [3,4]. These learning propensities can also evolve in response to changes in conditions. In experimental conditions where colour cues are reliable, but odours are not, *Drosophila* evolve strong visual learning propensities [5]. An animal’s learning abilities are therefore influenced by the relative reliability of the cues associated with key resources [6]. However, the context in which a cue is encountered can also have a role in shaping the reliability of that cue [7,8].

For many pollinators, foraging for flowers occurs in the context of temporal variation in resource profitability. Flowers tend to vary predictably in the availability of pollen and nectar [9]. Consequently, some specialist nectarivores use time as a contextual cue to modulate their foraging strategy [9]. For example, honey bees can consistently change their preference towards particular visual cues throughout the day [10,11], and some nectarivorous ants remember the time of day at which resources are most profitable [12,13]. However, the ability to learn time-dependent associations has only been formally demonstrated in a handful of invertebrates, including *Drosophila*, bees, and ants [11,14–21], and there is evidence that this ability can vary across species from the same family [12,13,22]. Hence, the prevalence of this ability, and the ecological factors that may account for its variability, are unclear.

*Heliconius* butterflies display a foraging behaviour not seen in the other 17,000+ described species of butterflies [23–25]. *Heliconius* actively collect and feed on pollen grains from a restricted range of plants that occur in low densities, but flower continuously [23,26]. This dietary adaptation provides an adult source of essential amino acids [24], which is plausibly linked to dramatically increased lifespan and reproductive longevity [27]. Pollen feeding is associated with a suite of derived foraging and cognitive adaptations not seen in other tropical butterflies, including fidelity to a local home-range [26,28], as well as temporally and spatially faithful ‘trap-lining’ behaviour, in which individual butterflies consistently visit particular flowers at specific times of day [26]. Foraging efficiency is reported to increase with experience, suggesting trap-lines are learnt and refined throughout an individual’s life [29]. Recent data also suggest wild *Heliconius* visit particular flower species in a manner that coincides with the maximal temporal availability of nectar and pollen rewards [30]. Since the timing of pollen release and nectar production varies across flowering plants [26,30], the time of day becomes a useful contextual cue to optimise foraging behaviour [26,30].

Given their derived foraging behaviour it has been hypothesised that *Heliconius* have evolved specific cognitive traits that support trap-lining behavior, including the ability to use the time of day as a contextual cue [26,31]. However, to our knowledge, time-dependent associative learning has not been reported in any Lepidoptera. In this study, we provide the first evidence that *Heliconius* butterflies can form time-dependent preferences for distinct flowers, and subsequently explore whether this ability is a derived trait in *Heliconius* associated with the evolution of pollen-feeding.

## 2. Materials and Methods

Experiments were performed at the Smithsonian Tropical Research Institute in Gamboa, Panama. We initially focused on *Heliconius hecale*, with secondary experiments in *Heliconius melpomene* and *Dryas iulia*, a closely related, non-pollen feeding Heliconiini. Individuals were labelled with unique IDs. Animal husbandry is described in the ESM.

### 2.1. Experimental set-up

Artificial feeders were constructed from foam sheets with an Eppendorf tube placed centrally. Yellow and purple were chosen as experimental colours as they were equally approached by naïve butterflies in our pilot experiments. During training, feeders contained either a 10% sugar water solution with 2.5 CCF per 50 ml of critical care formula (a surrogate for pollen; rewarded feeder), or a saturated quinine water solution (punished feeder). 12 artificial feeders of each colour were placed on a grid of 24, with randomised positions (Figure S1). Butterflies were presented with feeders for 2 hours in the morning (AM) (08:00-10:00) and 2 hours in the afternoon (PM) (15:00-17:00).

### 2.2. Experimental procedure

The experiment had four phases. 1) During pre-training, freshly eclosed butterflies were fed on white artificial feeders, in the AM and PM, for two days. 2) The naïve shift in colour preference based on time of day was recorded prior to training, using clean, empty feeders. Due to low feeding rates in the PM session, we split the initial preference test across two days. AM colour preference was recorded on day 1, and butterflies were food deprived in the PM. PM colour preference was performed on day 2, after food deprivation in the AM. 3) The training reward structure was split such that the purple feeders were rewarded in AM and yellow feeders rewarded in PM, or vice versa. This training phase lasted for 10 days. 4) During the final post-training preference test butterflies were presented with clean, empty feeders for one hour in the AM, followed by the reinforced AM feeders for one hour, and then clean, empty feeders for an hour in the PM. To determine whether butterflies were learning the order in which they encountered the reward, rather than the time of day, a proportion of butterflies had the order of their AM and PM trials reversed (see supplemental material). During trials, artificial feeders were filmed from above with a GoPro HERO 5 camera (Figure S1). Using this footage, we scored the number of feeding attempts made by each individual.

### 2.3. Training criterion

For an observed behaviour to be a consequence of learning an animal must experience the reward contingency scheme [32]. Individuals were significantly less active in AM than PM during training (*z* = −13.11, *n* = 41, p < 0.01, figure S1A). As a consequence, some individuals (*n* = 11) either did not attempt to feed from both feeders in AM or, more commonly, PM during training, or did not make any foraging attempts during a final test session, and were removed from further analyses. We analysed the remaining data in two ways. First, we ran models including all remaining individuals. Second, following previous learning studies [33–35], we established a training criterion. As we are interested in whether time-dependent memories are formed and can therefore guide behaviour in the absence of the reward, we identified individuals that correctly adjust their behaviour in AM and PM sessions during training with reinforced feeders. We then asked whether these individuals demonstrate evidence of learning by behaving in the same way when presented with unreinforced feeders in the post-training preference test. Our training criteria was that the majority (>50%) of feeding choices made by an individual in the final two training days were correct in both AM and PM.

### 2.4. Statistical Analysis

Data were analysed using generalized linear mixed models (GLMMs) in R [36]. We examined the influence of time on: (a) activity levels, measured as total foraging attempts, using GLMM with a Poisson distribution; (b) shifts in proportional colour preference when naïve, using GLMM with a binomial distribution; (c) shifts in proportional colour preference when trained, using GLMM with a binomial distribution, and presentation order in the final test included as a predictor. Across all models we included identity as a random effect. As individuals were trained and tested in groups of 8-13 within a single cage, where possible, a random effect of cage was also included to control for group level cage effects. We ensured all models fit their assumptions with the R package DHARMa [37] (see ESM).

## 3. Results

### 3.1. Heliconius *butterflies can learn time-dependent associations*

Across the *H. hecale* that experienced the full training set, there was no significant effect of the time of day on naïve colour preferences (z = 0.90, n = 30, p = 0.36), and no overall effect of time of day on trained colour preference (z = −1.846, n = 30, p = 0.06). However, there was considerable variation in behavior during training, and only a subset of individuals (n = 16) passed the training criterion (Figure S2). Prior to training, both butterflies that met the training criterion, and those that did not, showed no significant shift in colour preference throughout the day (z = 0.33, n = 16, p = 0.73 and z = 1.15, n = 14, p = 0.24 respectively).

Once trained, however, individuals that passed the training criteria showed a significant effect of the time of day on colour preference (z = −2.24, n = 16, p = 0.02, figure 1B). On average, the preference for AM rewarded colour decreased by 11% from AM to PM. The presentation order of the post-training preference test (AM first vs PM first) had no effect (z = 0.36, p = 0.71, n =16). Among individuals that did not meet the training criterion there was no shift in colour preference throughout the day after training (z = 1.05, n = 14, p = 0.29). Addition of a small sample of *H. melpomene* (n = 6) support and strengthen these results (see ESM).

**Figure 1:**
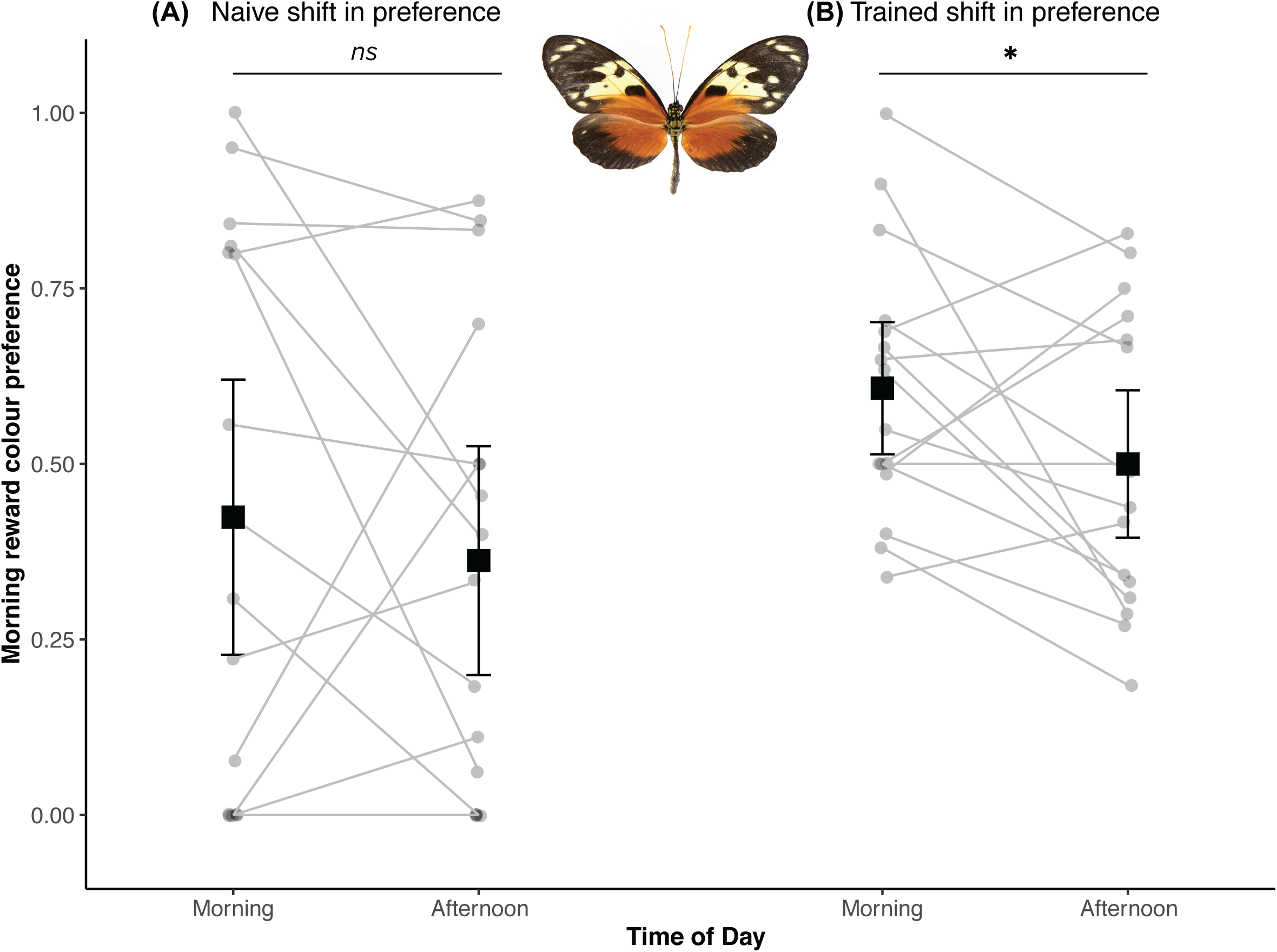
Data from the colour preference trials of *H. hecale* individuals who met the training criterion. Artificial feeders were new, empty, and unreinforced in both cases. (A) Naïve preferences of butterflies in the morning and afternoon. (B) Colour preferences of butterflies from (A) after training. Grey lines connect individuals across time periods. Data are presented as means ± SE. * indicates P <0.05

### 3.2. Evidence time-learning is common across Heliconiini

In a secondary experiment using *Dryas iulia*, a closely related genus within Heliconiini that does not pollen feed, 12 individuals experienced the full training set, with no overall response to training (z = 0.01, n = 12, p = 0.99). However, consistent with data from *H. hecale*, variation in the behaviour during training resulted in only a subset of individuals (n = 6) passing the training criterion. Among these individuals there was no significant effect of time of day on naïve preference (z = 1.67, *n =* 6, *p* = 0.09), but post-training there was a significant effect of time on colour preference, with preference for in the AM rewarded colour decreasing by 40% from AM to PM (Fig 2B, z = −9.334, n = 6, p < 0.001). Individuals that did not reach the training criterion show no effect of time before training (z =0.437, n= 6, p = 0.66), and show a significant shift in the incorrect direction post-training (z = 7.354, n = 6, p < 0.001).

**Figure 2:**
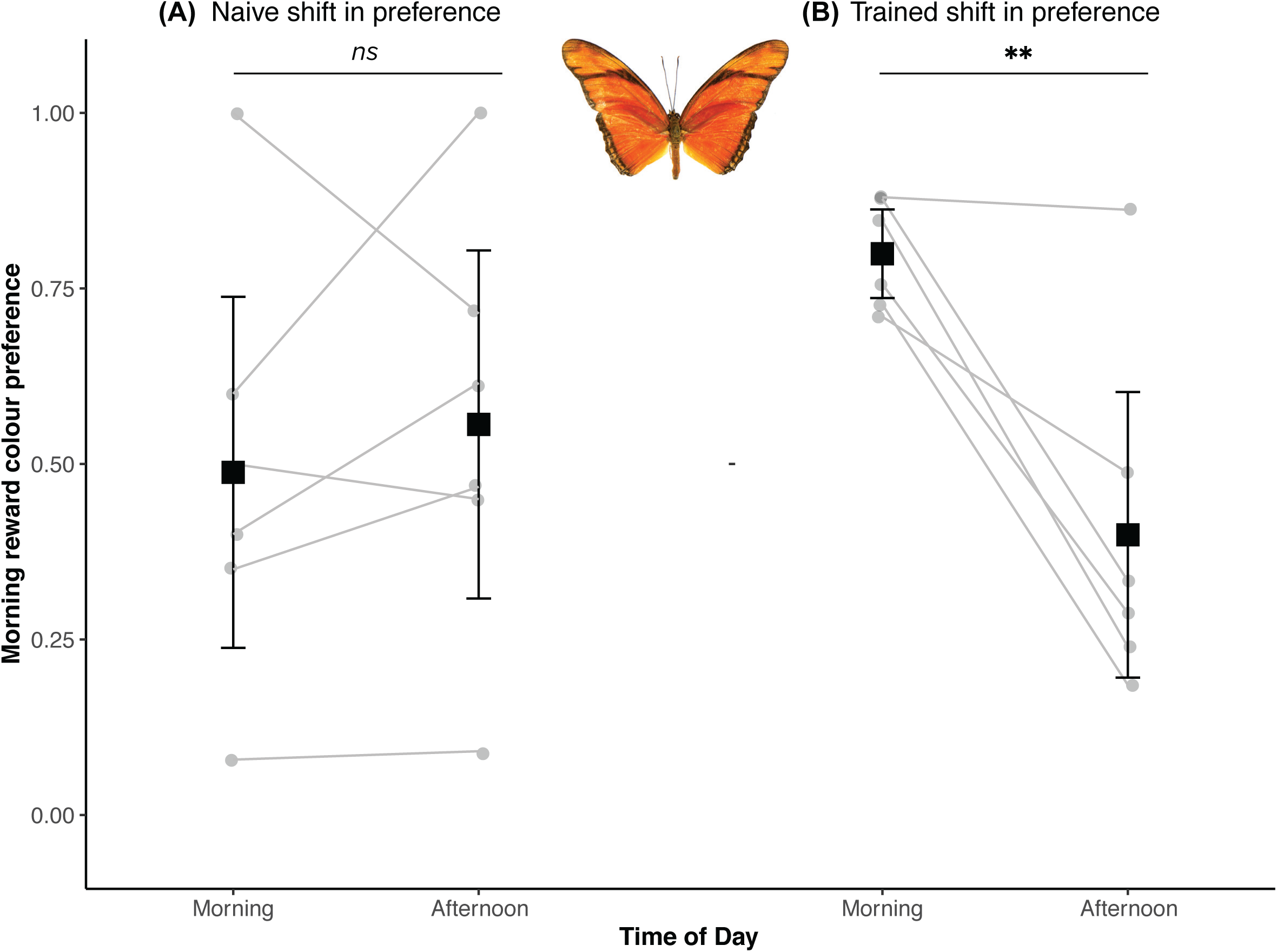
Data from the colour preference trials of *D. iulia* individuals who met the training criterion. Artificial feeders were new, empty, and unreinforced in both cases. (A) Naïve preferences of butterflies in the morning and afternoon. (B) Colour preferences of the butterflies from (A) after training. Grey lines connect individuals across time periods. Data are presented as means ± SE. ** indicates P<0.01

## 4. Discussion

In this experiment, we demonstrate that *Heliconius* butterflies can use time as a context for making foraging decisions. The observed shift in preference here is similar in magnitude to observed temporal variation in floral visits by wild *Heliconius* [30]. Time-dependent learning and trap-lining can occur via an ordinal or a circadian timing mechanism [38]. Given presentation order has no effect on our results, we find no support for ordinal learning as an explanation, indicating the possible presence of a circadian memory. While suggestive of a circadian mechanism, our data do not confirm the presence of an endogenous mechanism, as it is possible butterflies are responding to external cues that covary with the time of day (*e.g*. light levels, sun position). Nevertheless, the functional consequences of time learning by either mechanism are similar. To the best of our knowledge, these results provide the first experimental evidence of time-dependent learning in Lepidoptera.

The dietary innovation of pollen feeding in *Heliconius* butterflies has had major implications for their life history and ecology [28,39]. For most butterflies, reproductive output is constrained by the proteins collected during the larval stage [40]. Pollen feeding provides adult *Heliconius* with a reliable source of protein, leading to a pronounced delay of reproductive senescence [41]. Being able to efficiently exploit the sparsely distributed pollen-rich flowers is therefore critical to reproductive success in *Heliconius* [41]. Competition for pollen resources can be pronounced, and some *Heliconius* will forage early in the morning and actively defend flowers against other butterflies [23], suggesting selection may have favoured cognitive mechanisms that increase foraging efficiency. On this basis, it has been suggested that *Heliconius* may have acquired the ability to use time as a foraging cue in the context of pollen-feeding [26]. Indeed, our experiments confirm *Heliconius* can use time as a foraging cue. However, we also show that a non-pollen feeding relative, *Dryas iulia*, has the same capacity. This suggests that an ability to learn time-dependent associations did not evolve in response to selection for trapline foraging, and pre-dates the origin of pollen-feeding. Although our sample size for *Dryas* is smaller than for *Heliconius*, the proportion of individuals passing the training criterion and the pattern of results are highly consistent. While data on the foraging behavior of *Dryas* in the wild is limited, they have no known foraging specialisations that are not seen in other groups, and have lifespans typical for tropical species [42]. It is therefore reasonable to suggest that the ability to use time as a contextual foraging cue may be widespread across butterflies.

Previous work shows that time learning is prevalent among social Hymenoptera, where allocentric foraging provides an ecological context for using time cues in the context of a consistent foraging landscape [43–45]. *Heliconius* have converged on several foraging behaviours observed in these species, and also share dramatically expanded mushroom bodies, a region of the insect brain responsible for learning and memory [39]. Time-dependent memory acquisition is also reported in cockroaches [46], a third clade associated with mushroom body expansion [47]. This could be seen as indicating that the ecological challenges associated with learning foraging sites exert selective pressures favouring neuroanatomical elaboration supporting specialised cognitive abilities, like time learning [21,47–50]. However, our data on *Dryas* suggest that elaborated mushroom bodies are not necessary for the time learning abilities displayed in these taxa. This is further supported by the fact that *Drosophila*, which have substantially smaller mushroom bodies, can also learn time-dependent olfactory associations [14]. Therefore, the neural basis of integrating time information with foraging cues may be relatively simple. Integrating time and place memories may be more complex than forming these associations in isolation, as hypothesised in hummingbirds [48,51]. However, time learning is likely to be an important precursor for temporally and spatially faithful foraging. Hence, the pre-existence of this trait may have helped facilitate the evolution of trap-lining, and the transition to pollen-feeding in *Heliconius*.

The ability to form time-dependent associations may have wider ecological effects. If a pollinator has a time learning ability, sympatric plant species can coexist by sharing, rather than competing for, the same pollinator by temporally partitioning pollen or nectar rewards [28]. This effect may explain observed divergence in the nectar/pollen release schedules of *Psiguria* flowers, a preferred pollen resource for *Heliconius* [26]. A similar phenomenon is observed in bees and *Dalechampia* flowers, adding support to the idea that time learning abilities have implications for ecological diversity [28].

Overall, our results support the importance of temporal predictability in resources, rather than allocentric foraging or pollen-feeding, in promoting an ability for time learning. This is supported by the presence of time learning in both *Dyras* and *Heliconius*, and both pollinivorous and nectarivorous Hymenoptera. Moreover, time learning does not seem to be associated with an expansion of insect memory centres. Whether butterflies use internal or external cues to register time of day remains an open question for future study.

## Supporting information

Supplemental Results

## Ethics

This work was carried out under permission from the Ministerio del Ambiente, Panama (permit number: SE/AP-14-18).

## Data accessibility

Data and R scripts are available from the Dryad Digital Repository: (https://doi.org/10.5061/dryad.c59zw3r4m).

## Author’s contributions

MWT, FJY, SHM conceived and designed the experiment. MWT collected and analysed the data, and wrote the paper, with practical support and input from FJY and WOM, under the supervision of SHM.

## Acknowledgements

We thank Lina Gabriela, Laura Hebberecht-Lopez, Oscar Paneso and Cruz Batista Saez for support at the insectaries. We also thank the Reader and Guigueno labs (McGill), EBaB lab (Bristol), and the Butterfly Ecology and Evolution Research group (Smithsonian) for comments on the manuscript. We thank Sebastian Mena for the images of butterflies for the figures. Finally, we are grateful to the Ministerio del Ambiente, Panama for collection permits and facilitating this work.

