## Supplemental Results for "Heliconiini butterflies can learn time-dependent reward associations"

**Contents:**

In the ESM we provide additional methodological detail for clarity, and additional results supporting our conclusions. Some information may be repeated from the main text to make the supplementary material easier to follow.

1. **Extended materials and methods**
   1.1. Animal husbandry, page 2
   1.2. Experimental procedure with graphical summary, page 3
   1.3. *Dryas* experiments, page 5
   1.4. Inclusion of the training criterion, page 6
   1.5. Statistical analysis, page 10
2. **Supplemental results**
   2.1. Individuals passing the training criteria, page 11
   2.2. Effects of including data from *H. melpomene*, page 13
   2.3. Results grouping all data, page 14
3. **Supplemental references**

**1. Extended materials and methods**

**1.1 Animal husbandry**

All butterfly larvae were reared from outbred stocks of wild caught individuals in ambient conditions. *H. hecale* were reared on a mixture of *P. platyloba* and *P. vitifolia*. *H. melpomene* were reared on *P. triloba* and *P*. *vitifolia*. *Dryas iulia* larvae were reared on *P. biflora.* Larvae were reared in mesh containers and transferred to a larger mesh contained upon pupation until the adult butterflies eclosed.

Adult butterflies were labelled with unique IDs, using numbers written in ink on the ventral side of the forewings and dorsal side of the hindwings, which were clearly identifiable when seen on recorded footage.  2x2x2m stock and experimental cages were made of a mesh net that allowed butterflies to live in ambient conditions and natural light conditions (Figure S1). Cages also included *Psychotria elata* as a roosting site, but with all flowers and flower buds removed.

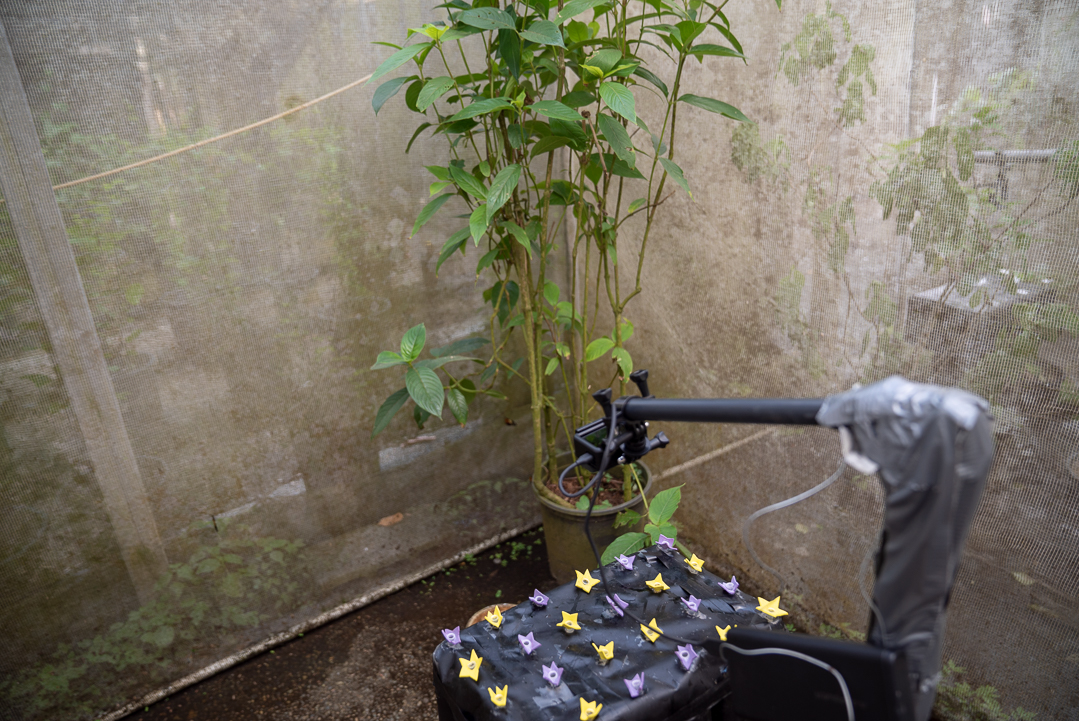

Figure S1: Photograph of the recording set up in an experimental cage.

**1.2 Experimental procedure with graphical summary**

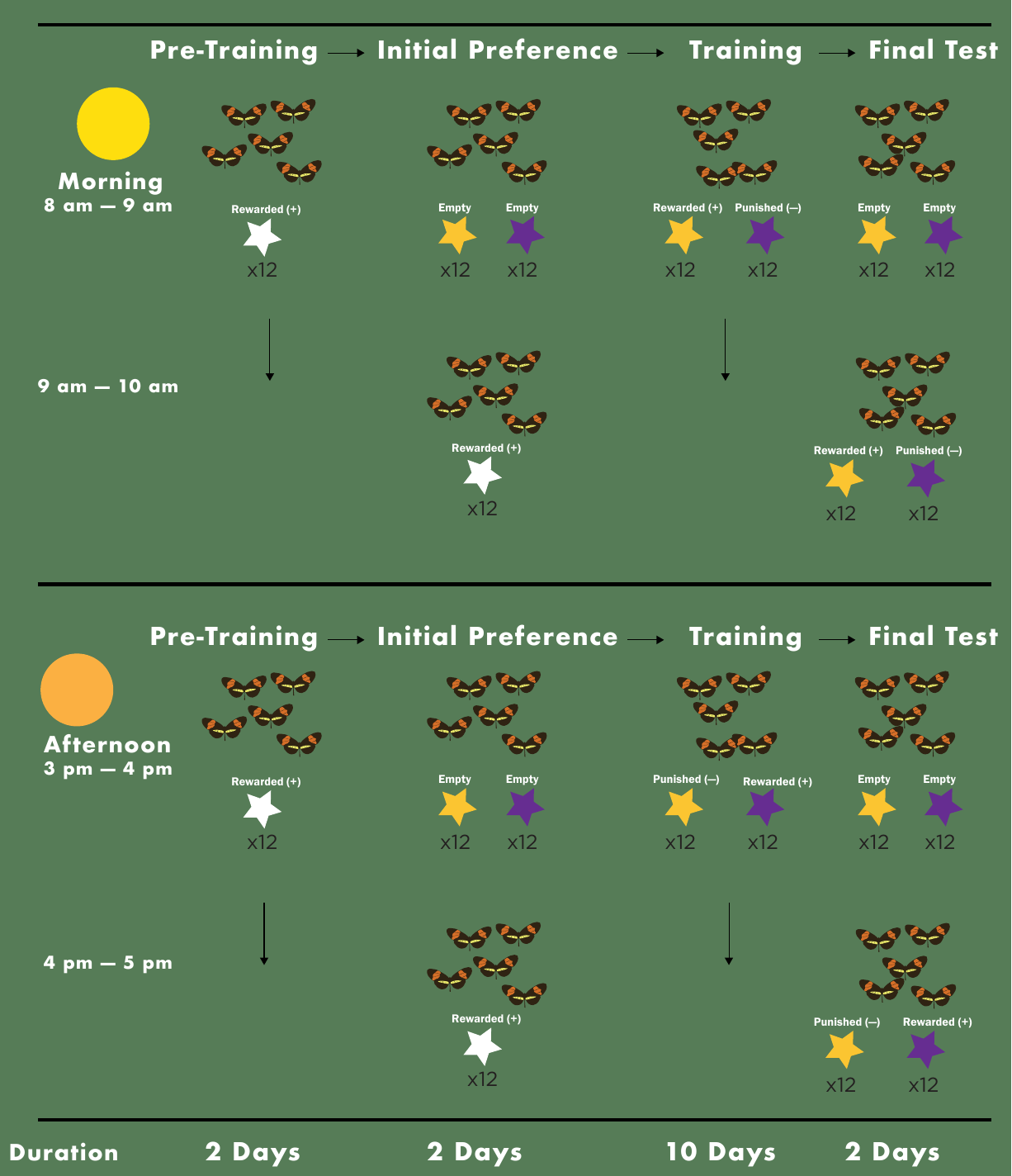

Figure S2: Graphical summary of the experiment. Showing the procedure for a single group of butterflies. Arrows indicate the condition from the previous hour carries over into the next hour. In a group with the final test presentation order reversed the morning session would have no feeders in AM followed by a PM session test, PM session reinforced feeders, and then an AM session test the next day.

The experiment had four phases.

**1) Pre-training**: One day after eclosion individuals were transferred to an experimental cage for pre-training. Butterflies were presented with feeders for 2 hours in the morning (AM) (08:00-10:00) and 2 hours in the afternoon (PM) (15:00-17:00). During pre-training, butterflies were fed on white feeders in both the morning and afternoon for two days, to accustomise them to the use of artificial feeders.

**2) Initial preference:** The colour preferences for the yellow and purple feeders were recorded prior to training, using clean, empty feeders to ensure the response was visually determined. Due to low feeding rates in the PM during pilot experiments, we split the initial preference test across two days. We gave the butterflies an AM preference test on day 1, and then food deprived them in the PM. On day 2 we deprived them of food in the AM and gave them a preference test in the PM. After recording naïve preferences, the training phase began.

**3) Training:** The training reward structure was split such that a butterfly either experienced purple feeders as being rewarded in the AM, and yellow feeders as being rewarded in the PM, or vice versa. The training phase lasted for 10 days before individuals moved on to the final preference tests.

**4) Final preference**: For the final preference tests the butterflies were presented with clean, empty feeders for one hour in the AM, followed by the reinforced morning reward feeders for one hour, and then clean, empty feeders for an hour in the afternoon. To determine whether butterflies were using the order in which they encountered the reward rather than the time of day, a proportion of butterflies had their AM and PM trials reversed. This reversed group received empty feeders for an hour in the PM, followed by reinforced feeders for an hour, and then empty feeders for an hour in the AM of the next day.

For an observed behaviour to be a consequence of learning an animal must actually experience the reward contingency scheme [1]. Individuals that did not experience both colours in both time points at least once during the whole training period (*e.g.* never making a correct choice in the PM during all of training) would not have the opportunity to learn time-dependent associations. *H.* *hecale* and *H. melpomene* individuals were significantly less active in the PM than in the AM throughout training, with *H*. *hecale* making on average 24 less foraging attempts in the afternoon (*z* = -13.11, *n* = 41, p < 0.01, figure S3A) and *H. melpomene* making on average 44 less foraging attempts in the afternoon (*z* = -13.31 *n* = 10, p < 0.01, figure S3B). *D. iulia* individuals made on average 15 more foraging attempts in the afternoon (*z* = 6.634, *n* = 19, p < 0.01, Figure S3C). As a consequence, some individuals either did not attempt to feed from both feeders in AM or PM during training or did not make *any* feeding attempts during a final test session, and were removed from further analyses.

In total, 41 *H. hecale* and 10 *H. melpomene* individuals went through the experiment. Due to space restrictions, individuals were trained and tested in groups of 8-13 within a single flight cage. In total 5 cages were used for *H. hecale*; 3 had yellow rewarded in the AM and purple rewarded in the PM, while 2 had purple rewarded in the AM and yellow rewarded in the PM and one of each reward treatment had the final preference presentation order reversed. For *H. melpomene* there was only one cage of individuals and it had yellow rewarded in the AM and purple in the PM with an unreversed presentation order (AM then PM). 39 out of 51 individuals made feeding attempts at both colours across both time periods, the remaining 12 were removed from analysis. Additionally, 3 individuals did not make any choices during the final test and were removed from analysis, leaving 36 individuals (30 *H. hecale* and 6 *H. melpomene*).

**1.3 *Dryas* experiments**

In a second experiment, we tested *Dryas iulia*, a closely related butterfly, to test if the ability to learn associations was limited to *Heliconius*. 13 out of 19 *D. iulia* made both choices in both time periods during training, the remaining 5 were removed from further analyses. An additional individual was removed after making no feeding attempts in the final test. One cage had purple rewarded in the AM and yellow rewarded in the PM with an unreversed final test order (group 1), while the second cage had yellow rewarded in the AM and purple rewarded in the PM with a reversed final test order (group 2). In this sample only *Dryas* from group 2 met the training criterion (*n* = 6). However, since the order of the final test has no bearing on how an individual behaves during the training trials, and test order has no effect in *H. hecale*, this is unlikely to be problematic. Moreover, the proportion of individuals passing the training criterion is statistically consistent with the *H. hecale* and *H. melpomene* results; of the individuals that were included in the analysis, 50% of *D. iulia,* 50% of *H. melpomene*, and 53% of *H. hecale* pass the training criterion (3 sample test of equal proportions, X^2^ = 0.05, p = 0.97, figures S8-S10).

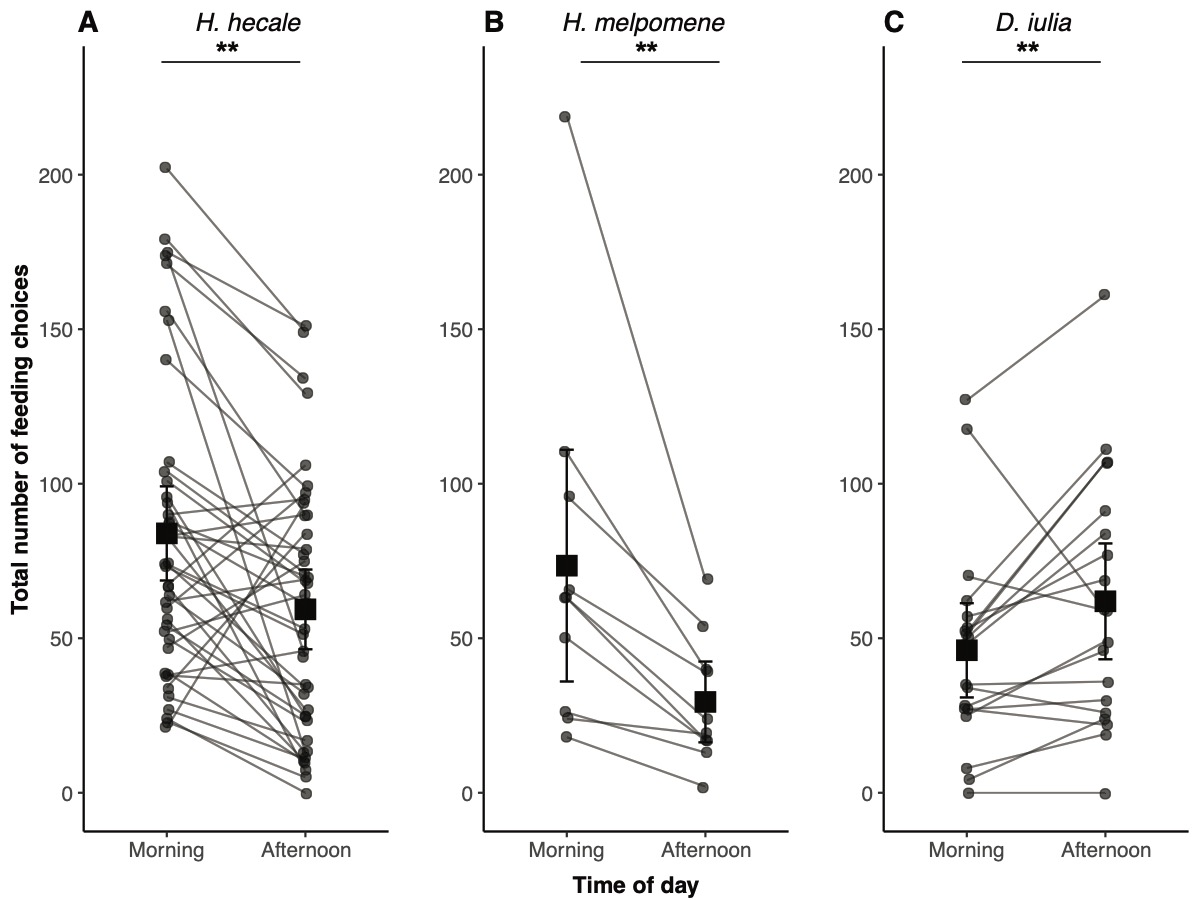

Figure S3: Total feeding choices made by all butterflies throughout training. Feeding activity was significantly lower in the afternoon for the two Heliconius species, while Dryas activity increased in the PM . Grey lines connect individual butterflies across time periods. Data are presented as means ± SE

**1.4 Inclusion of the training criterion**

Following previous learning studies [2–6], we established a training criterion. Several studies interested in learned behaviour restrict statistical analyses to individuals which show a response during training, as this indicates they are using the relevant cue to guide their behaviour. Note, that since this may be a short-term reaction to the reward and associated stimuli, it does not presume learning has occurred. Since we are interested whether individuals had were adjusting their behaviour in both AM and PM sessions during training were forming stable time-dependent memories of the feeder rewards, we set a similar criterion. Our training criterion was that the majority of feeding choices (greater than 50%) made by a butterfly were correct in their last 2 days of training, in both the AM and PM sessions. Individuals that were not making a majority of correct choices were deemed to be unresponsive to the contextual cue, and therefore not expected to have learned the association. Post-hoc support for this is provided by evidence that individuals that were indeed responding correctly during training displayed evidence of learning in the final test with unreinforced feeders, whereas those that were not yet making a majority of correct responses did not display evidence of learning in the case of *H. hecale* and *H. melpomene,* or had a weak but significant shift in preference in the *incorrect* direction as seen in *D. iulia* (see main text and figures S4-6).

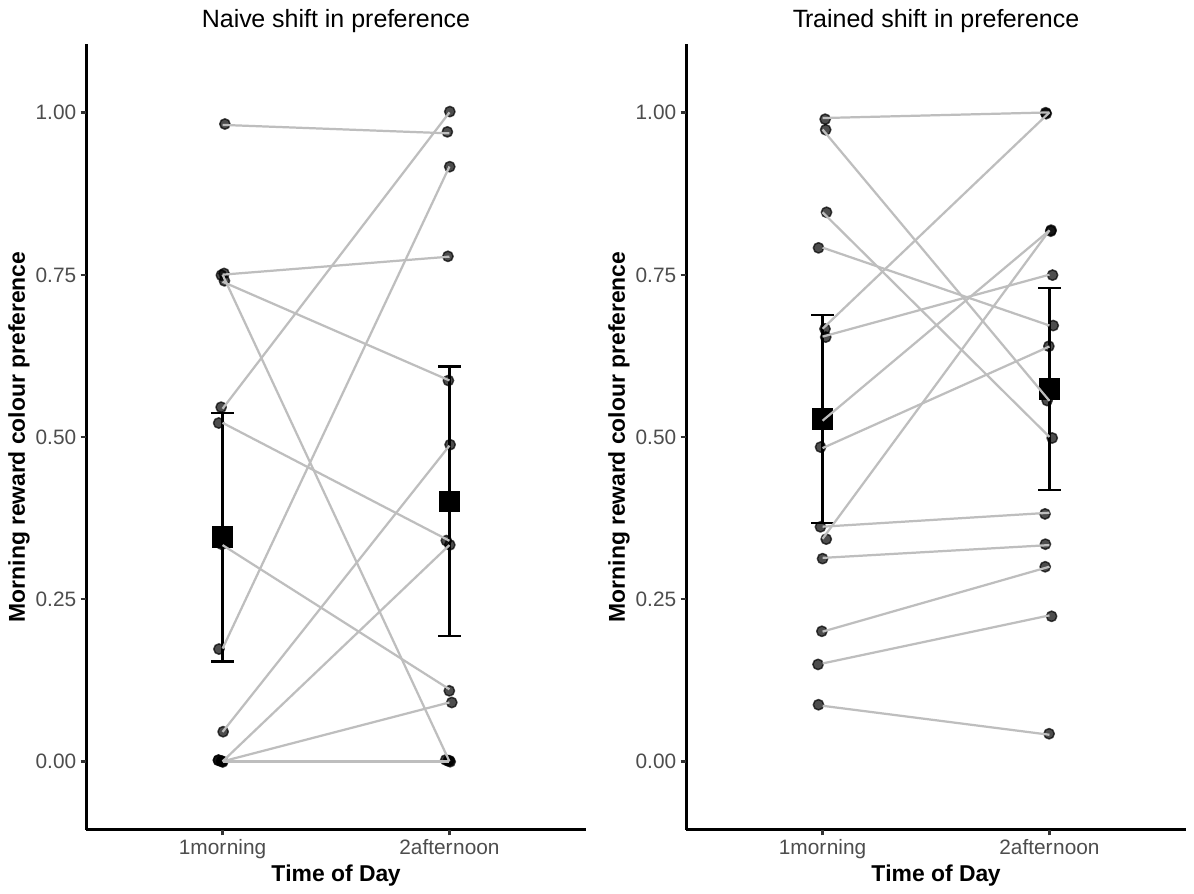

Figure S4: Shift in the proportional colour preference for the AM reward colour for H. hecale individuals that fail the training criterion.

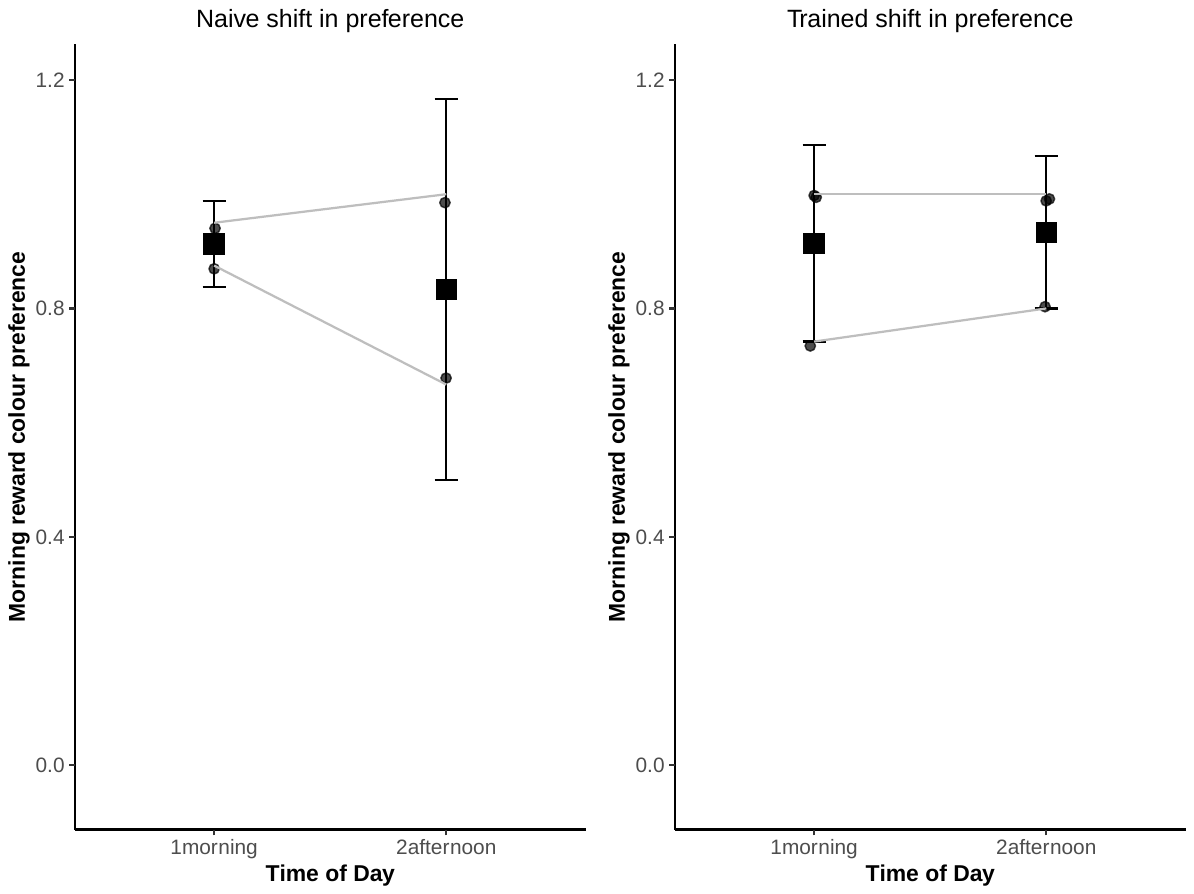

Figure S5: Shift in the proportional colour preference for the AM reward colour for H. melpomene individuals that fail the training criterion.

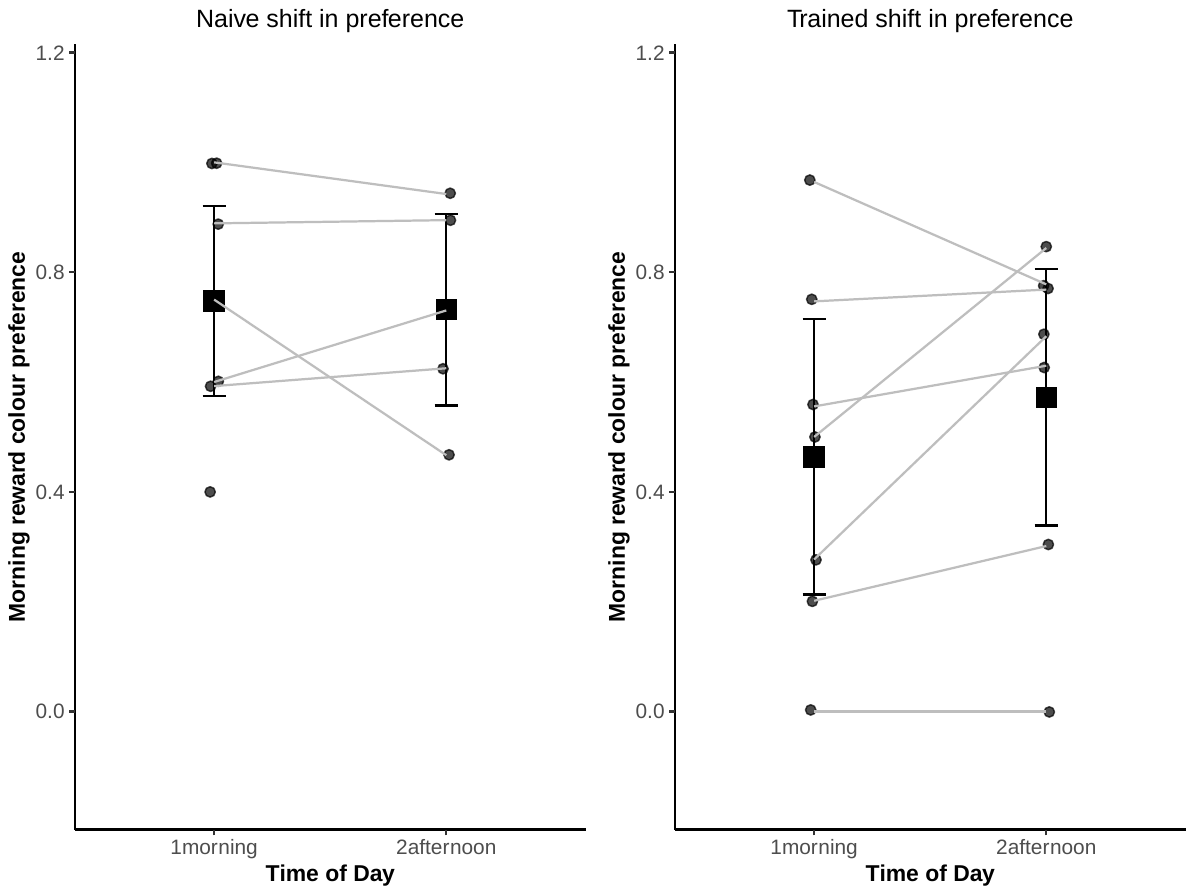

Figure S6: Shift in the proportional colour preference for the AM reward colour for D. iulia individuals that fail the training criterion.

We also explored why these individuals did not respond to the conditions in the training regime by examining the effects of the distribution of an individual’s colour preferences on their likelihood to pass the training criteria. Individuals that failed the training criterion tended to be more ‘inflexible’ in their choices during training. Analysing the shift in choices over training shows that individuals that failed the training criterion were more likely to consistently feed from one colour. This was quantified by the absolute difference between the proportion of choices for the morning reward colour in the morning session and the proportion of morning reward colour choices in the afternoon session over all choices made during training (figure S4). In figure S7 a value of 0 would indicate an individual that had a complete morning reward colour bias both AM and PM all throughout training, which would mean that this individual was staying fixed in their colour choice throughout training. The further from zero, the more ‘flexible’ the colour choices of an individual are throughout the two time periods. Lower values indicate less flexibility. Those that failed criterion had significantly lower shifts in morning reward colour preference (quasibinomial GLM, n = 50, t = 5.33, p < 0.001).

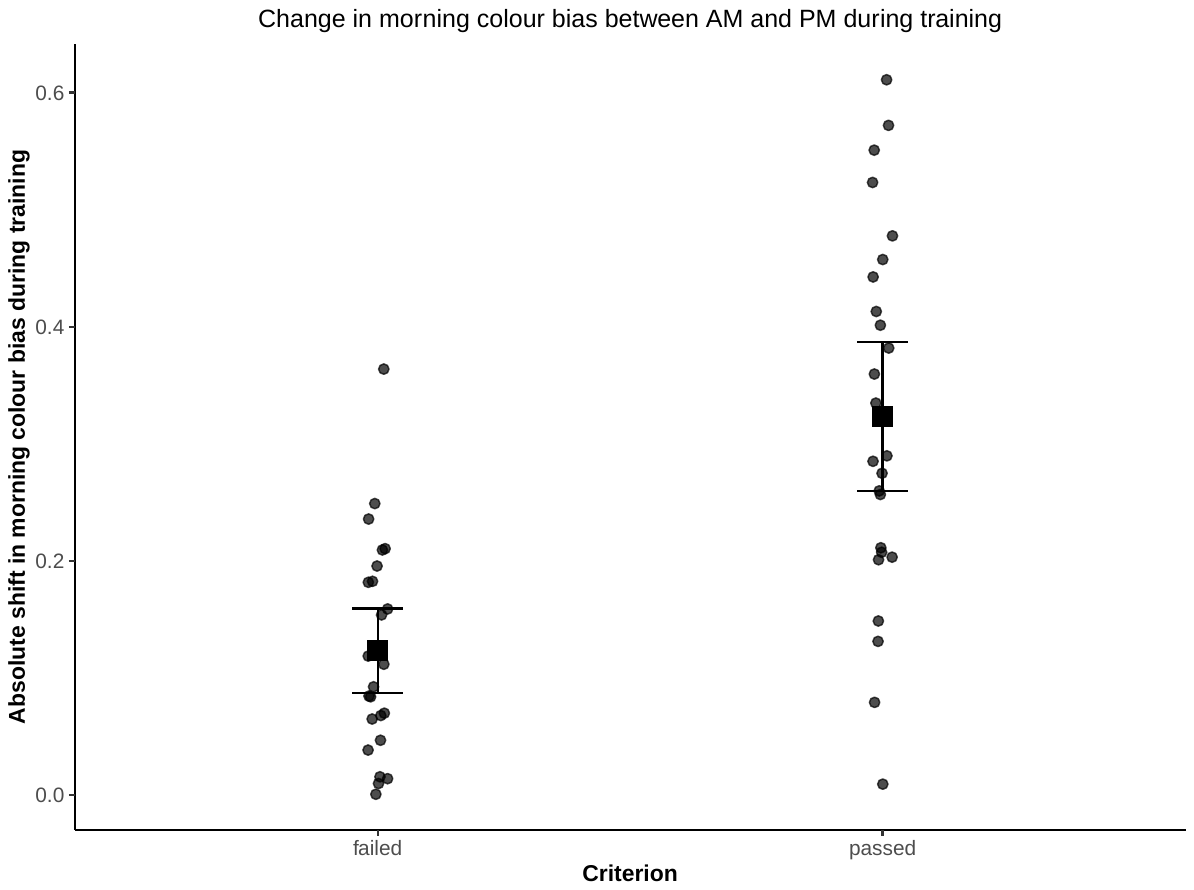

Figure S7: Choice flexibility during all of training for butterflies that passed or failed the training criterion. Butterflies that failed the training criterion were significantly more rigid in their colour preferences (lower shift in morning colour bias between AM and PM) than those butterflies which passed, indicating the vast majority of feeding attempts were made to one colour feeder regardless of time session.

**1.5 Statistical analysis**

Data were analyzed using generalized linear mixed models (GLMMs) in R using the lme4 package [7] with identity as a random effect due to repeated measures, as described in the main text. Where possible a random effect of cage was also included to control for group level cage effects. For models where individuals are split by meeting or failing the training criterion the number of individuals per cage becomes highly uneven leading to model convergence issues. For these models we dropped the cage random effect term and kept the random effect of individual ID. To ensure all models fit their assumptions models were checked for overdispersion, underdispersion, zero-inflation, and heteroscedasticity with the R package DHARMa [8]. The full R scripts with the model specifications and the commands to subset the individuals are included along with the full datasheet deposited in the Dryad digital repository (DOI: XXX).

**2. Supplemental results**

**2.1 Individuals passing the training criteria**

In the following plots we display the mean proportion of correct choices in the last two days of training for each individual in both time periods. Each species is displayed in a different column (indicated by numbered boxed). An individual’s points for both time periods must be above the dashed line to have passed the training criterion. Failed time points/individuals are indicated in red.

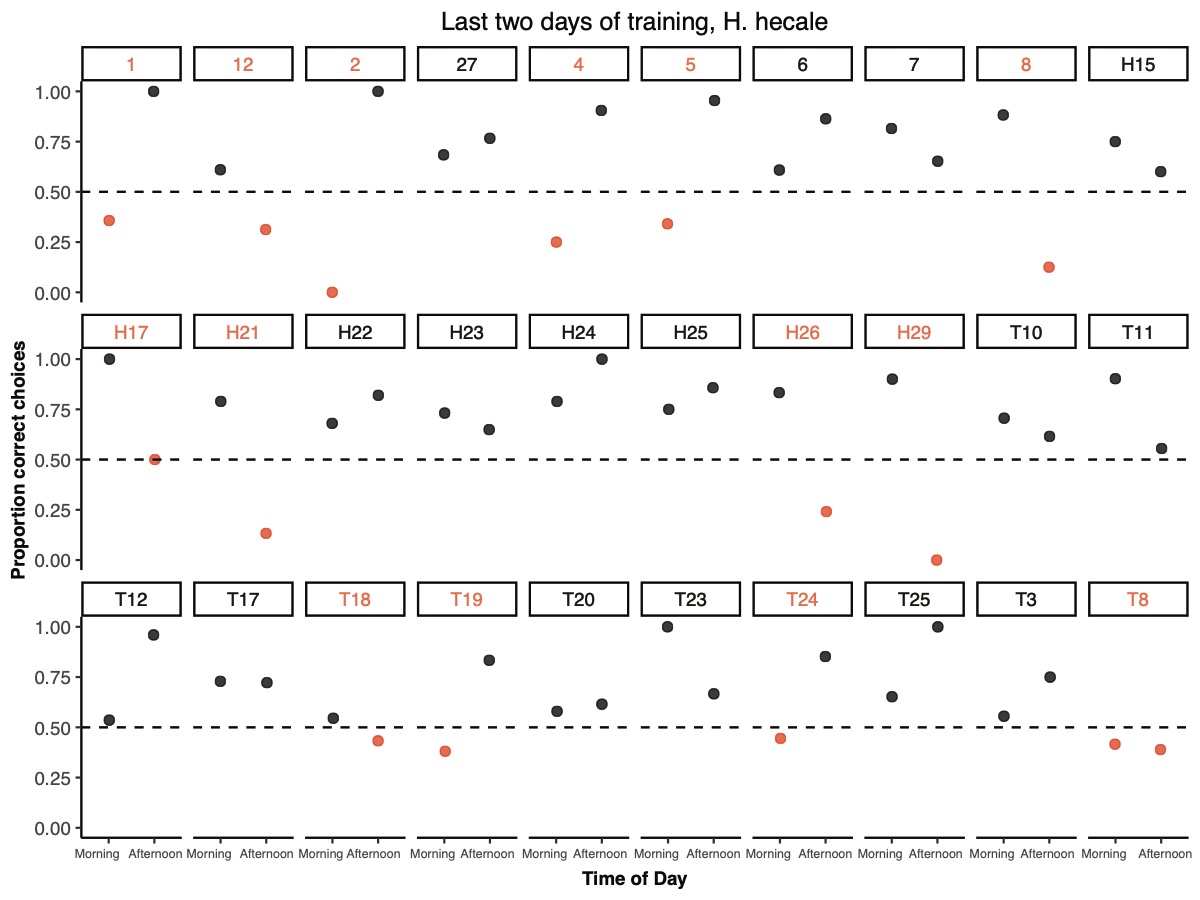

Figure S8: The proportion of correct choices made by H. hecale individuals during the two days of training prior to the unreinforced final preference test. Individual IDs are highlighted red to indicate that in at least one of two time periods they were below the training criterion of greater than 50% correct choices

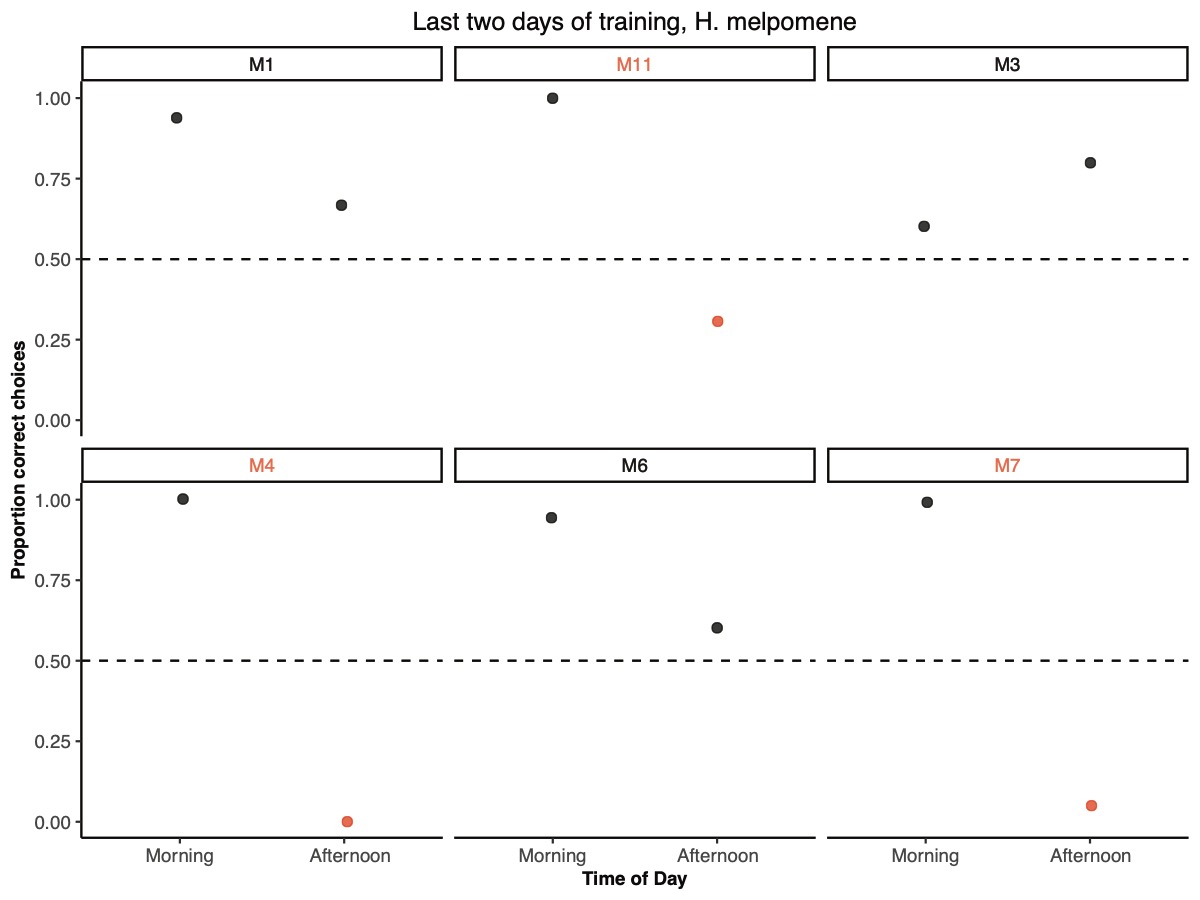

Figure S9: The proportion of correct choices made by H. melpomene individuals during the two days of training prior to the unreinforced final preference test. Individual IDs are highlighted red to indicate that in at least one of two time periods they were below the training criterion of greater than 50% correct choices.

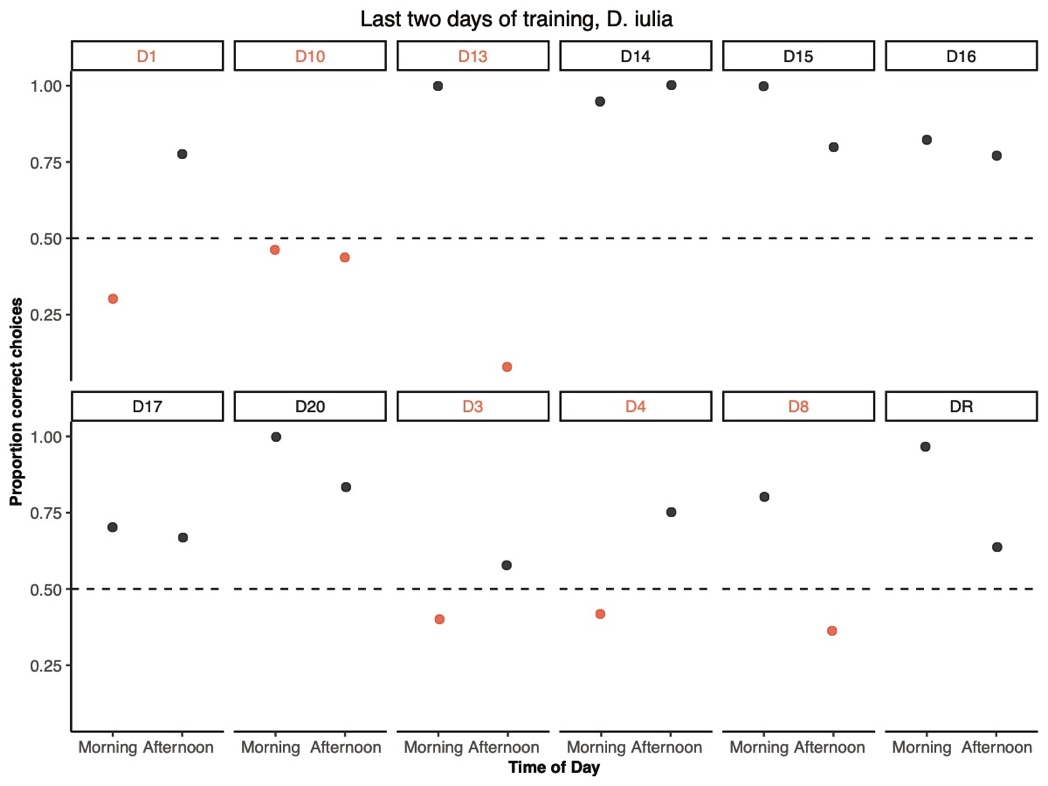

Figure S10: The proportion of correct choices made by D. iulia individuals during the two days of training prior to the unreinforced final preference test. Individual IDs are highlighted red to indicate that in at least one of two time periods they were below the training criterion of greater than 50% correct choices.

**2.2 Effects of including data from *H. melpomene***

*H. melpomene* were not included in the main text due to the small sample size and having only one final test presentation condition. However, the results in *H. melpomene* support and strengthen the results we find in *H. hecale.* Out of the 6 *H. melpomene* that were active during training and the final test, 3 passed the training criterion. These individuals significantly decreased their preference for the morning reward colour in the afternoon after training [z = -4.234, p < 0.001]. Including *H. melpomene* with the *H. hecale* dataset does not qualitatively change the results; individuals still display a significant decrease in preference for the AM reward colour in the PM (z = -3.578, n = 19, p< 0.001) and the effect size increases from an 11% shift in preference to a 17% shift in preference with no effect of final presentation (p = 0.22).

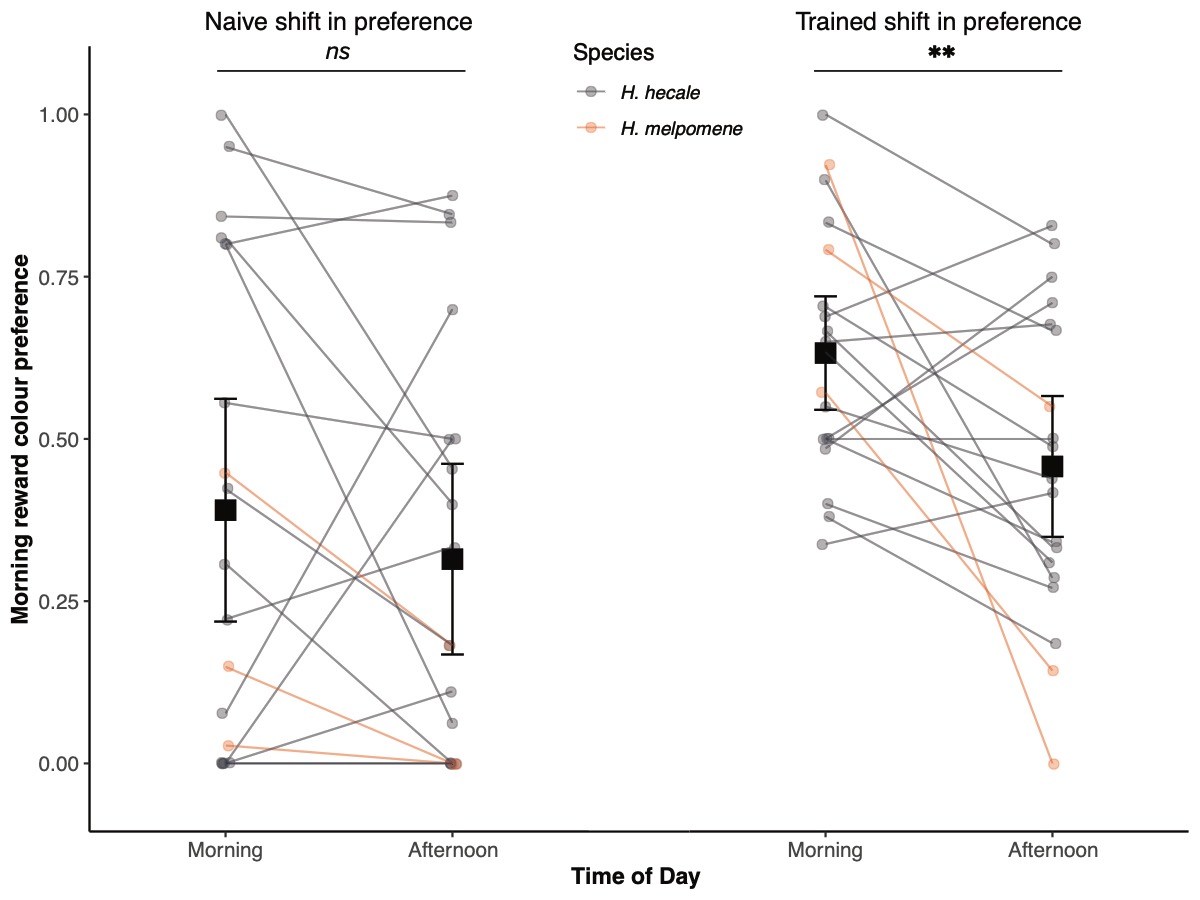

Figure S11: Shift in the proportional preference for the AM reward colour for individuals which pass the training criterion with H. melpomene individuals included in red. The preference for the AM reward colour significantly decreases in the PM.

**2.3 Results grouping all data**

Combining the data from all three species does not qualitatively change the results. There remains a significant effect of time of day on AM colour preference with the preference decreasing in the PM. However, there is an interaction effect with time of day and final presentation due to the disproportionately large effect size of the *Dryas* group, which was composed mainly of individuals in the reversed final presentation group. The shift in preference is therefore stronger in the group with the reversed final test presentation but the shift in preference is in the same direction for both treatments (figure S12). Since we were interested in whether the final presentation order would completely reverse the effect, leading to an increase for the morning reward colour in the afternoon, which would suggest ordinal learning, we drop the final presentation term when reporting the results for the grouped data. Across all individuals the preference for the AM reward colour significantly decreased in the PM (z = -8.611, n = 25, p < 0.001). On average the preference for the AM reward colour decreases by 23% in the afternoon. Therefore, the major conclusions throughout remain the same whether species are considered separately or grouped together.

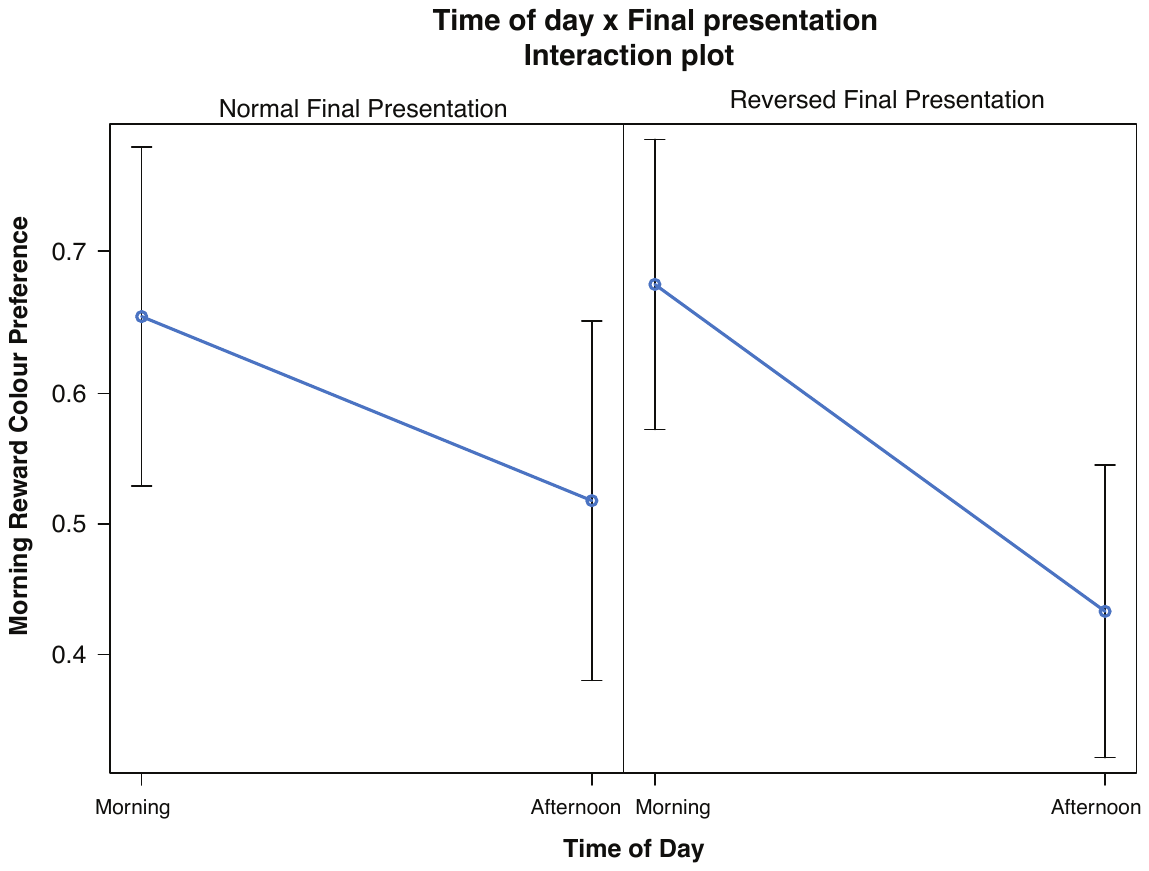

Figure S12: A visualisation of predictions of the GLMM on the final test data with individuals that passed the training criterion from all three species included. Both groups decrease their preference for the AM reward colour in the morning. There is an interaction effect where the individuals that had the final presentation reversed decreased their AM colour preference even more than those whom received a normal presentation.

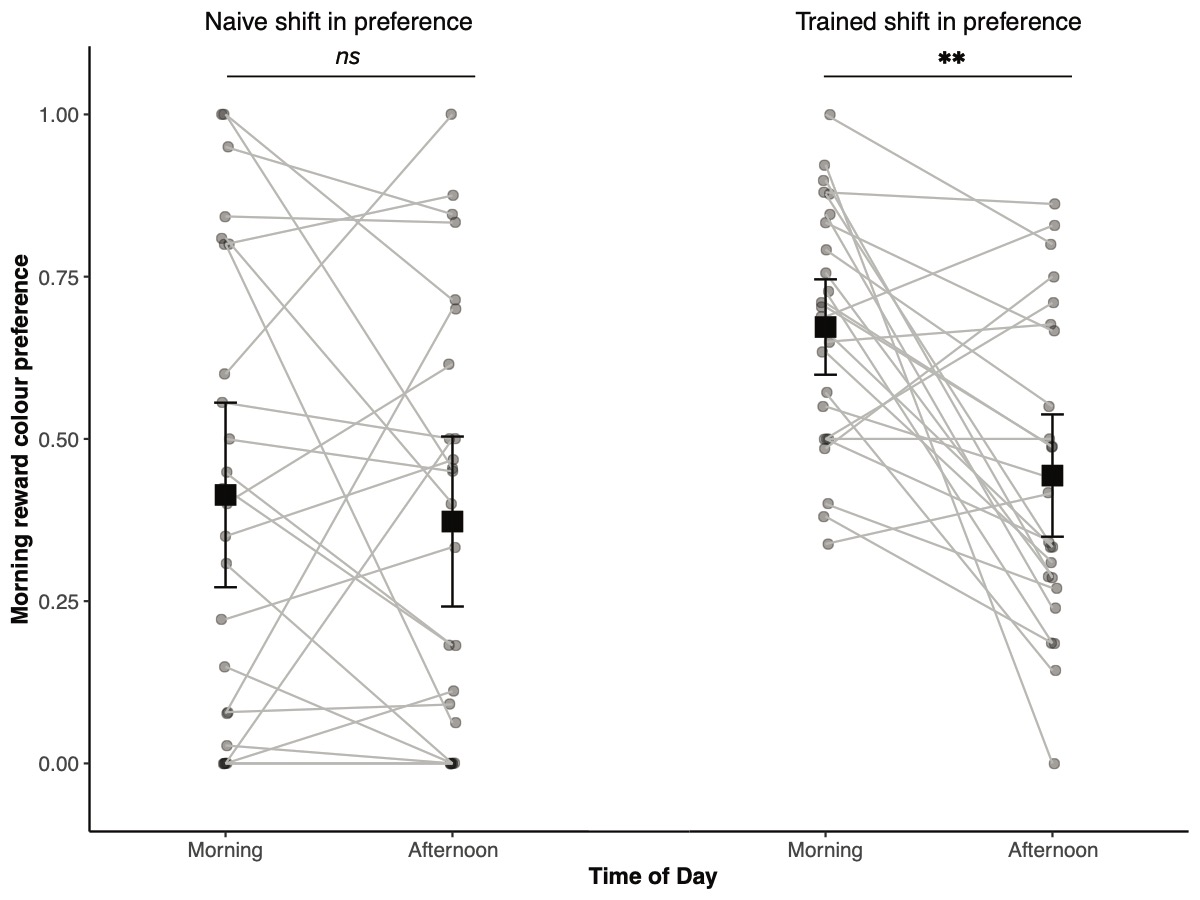

Figure S13: Shift in the proportional preference for the AM reward colour for individuals which pass the training criterion with all three species included.
